# Locational memory of macrovessel vascular cells is transcriptionally imprinted

**DOI:** 10.1101/2021.10.20.465092

**Authors:** Talitha C.F. Spanjersberg, Loes A. Oosterhoff, Hedwig S. Kruitwagen, Noortje A.M. van den Dungen, Magdalena Harakalova, Michal Mokry, Bart Spee, Frank G. van Steenbeek

**Affiliations:** Department of Clinical Sciences, Faculty of Veterinary Medicine, Utrecht University, Yalelaan 104, Utrecht, the Netherlands; Regenerative Medicine Centre Utrecht, University Medical Center Utrecht, Utrecht University, Uppsalalaan 8, Utrecht, The Netherlands; Department of Cardiology, Division Heart and Lungs, University Medical Center Utrecht, Utrecht University, Heidelberglaan 100, Utrecht, the Netherlands

**Author notes:** **Correspondence:** Frank G. van Steenbeek.

**Keywords:** genetic profiling, vascular heterogeneity, whole-transcriptome sequencing, weighted gene co-expression network analysis, locational memory

## Abstract

The locational predisposition of vascular pathologies illustrates the need for a better insight into vascular heterogeneity. To investigate the transcriptomic basis of angiodiversity, we isolated and analyzed transcriptomes from endothelial cells and vascular smooth muscle cells from nine different adult canine macrovessels: the aorta, coronary artery, vena cava, portal vein, femoral artery, femoral vein, saphenous vein, pulmonary vein, and pulmonary artery. We identified both reported and novel expression patterns defining specialized adult blood vessels. Our findings also show that adult vascular cells in culture express a remarkably high number of transcription factors crucial to organ development in the embryo. The persistent expression of these genes in culture indicates that these genes are not regulated by the flow or surrounding cell types but are rather fixed in the molecular memory. Therefore, our findings prompt the re-thinking of the extrapolation of results from single-origin endothelial cell systems.

## 1 Introduction

Various types of vascular disorders with spatial predilection, such as atherosclerosis, deep vein thrombosis, and pulmonary artery embolism, are known to be influenced by vascular heterogeneity.^1^ Nowadays, vascular specialization on a molecular level is being identified with increased specificity for different vessels under the emergence of large-scale transcriptome characterization technologies.^2, 3^ Although many studies focus on endothelial cells (ECs), other supporting cell types such as vascular smooth muscle cells (VSMCs) also play a role in vascular pathologies.^4^ To investigate how VSMCs compare to ECs in vascular heterogeneity, combining data from these cell types originating from a variety of vessels is needed. Obtaining such data, however, is hampered by the limited variety of vascular cell types that can be isolated from humans due to legal and ethical constraints. Therefore, large animals mimicking human physiology pose an interesting alternative tissue source, particularly considering the increasing demand for suitable animal models for the advancement of vascular tissue engineering.^5^ In an extension of our previous study, we used our recently published isolation protocol to isolate and culture both ECs and VSMCs from nine different canine macrovessels.^6^ Although endothelial heterogeneity has been reported to disappear after several passages, accumulating evidence demonstrates that some organ-specific expression patterns persist *in vitro*.^7–9^ These epigenetic features may be the foundation of vascular heterogeneity as they may explain the inadequate adaptations to hemodynamic forces in some vessels but not in others. For instance, back in the 70s, Wesly et al. transplanted a canine jugular vein segment into the arterial circuit of the carotid artery. Instead of acquiring arterial properties, the tissue response of the venous grafts was reflected by intimal proliferation, a common consequence of vascular injury.^10^ Nowadays, venous graft failure is still a frequent complication of bypass surgery, illustrating the need for insight into vascular heterogeneity.^11^

Studying the expression pattern of cultured cells from different anatomic locations may reveal an origin-dependent response to vascular injury. VSMCs are known to originate from various sources that partially may influence the disease susceptibility of vessels. For example, VSMCs in the aortic arch, embryonically derived from cardiac neural crest cells, are more prone to pathological calcification than VSMCs in the descending aorta originating from the mesoderm.^12^ These differences may persist *in vitro* even though cultured VSMCs are known to switch mainly to a proliferative synthetic phenotype. This switch from the typical quiescent contractile phenotype to a proliferative synthetic phenotype, accompanied by increased migration and extracellular matrix synthesis, is also an eminent mechanism of the response to injury.^13, 14^

The purpose of this study is to compare expression profiles of cultured ECs and VSMCs originating from various anatomical locations. A fundamental question relevant to vascular diseases and their complications is whether differential expression profiles *in vitro* can be linked to vascular functionality and disease susceptibility. Therefore, we generated a combined transcriptomic atlas for these nine vessel types and we correlated groups of co-expressed genes to vessel origin and cell type. Our observations extend knowledge to the fragmentary characterization of vascular subtypes and, therefore, assert the significance of using relevant cell types in cardiovascular research.

## 2 Results

### RNA sequencing identifies differentially expressed genes between vascular cells from different macrovessels

Recently we have analyzed the transcriptomes of canine ECs and VSMCs originating from the aorta, vena cava, and portal vein obtained by a novel isolation technique.^6, 15^ To further test the hypothesis that transcription patterns of vascular cells are origin dependent, we performed RNA-sequencing of cultured ECs (*n*=26) and VSMCs (*n*=27) originating from nine different macrovessels isolated from healthy dogs (*n*=4, **Fig. 1a**). Three samples were used from each cell and vessel type, except for coronary artery ECs, whereas only two samples of this group had sufficient viable cells. A total of 18,374 genes were detected at least once, and, after filtering for detection threshold, the dataset was brought back to 12,596 genes.

**Figure 1.**
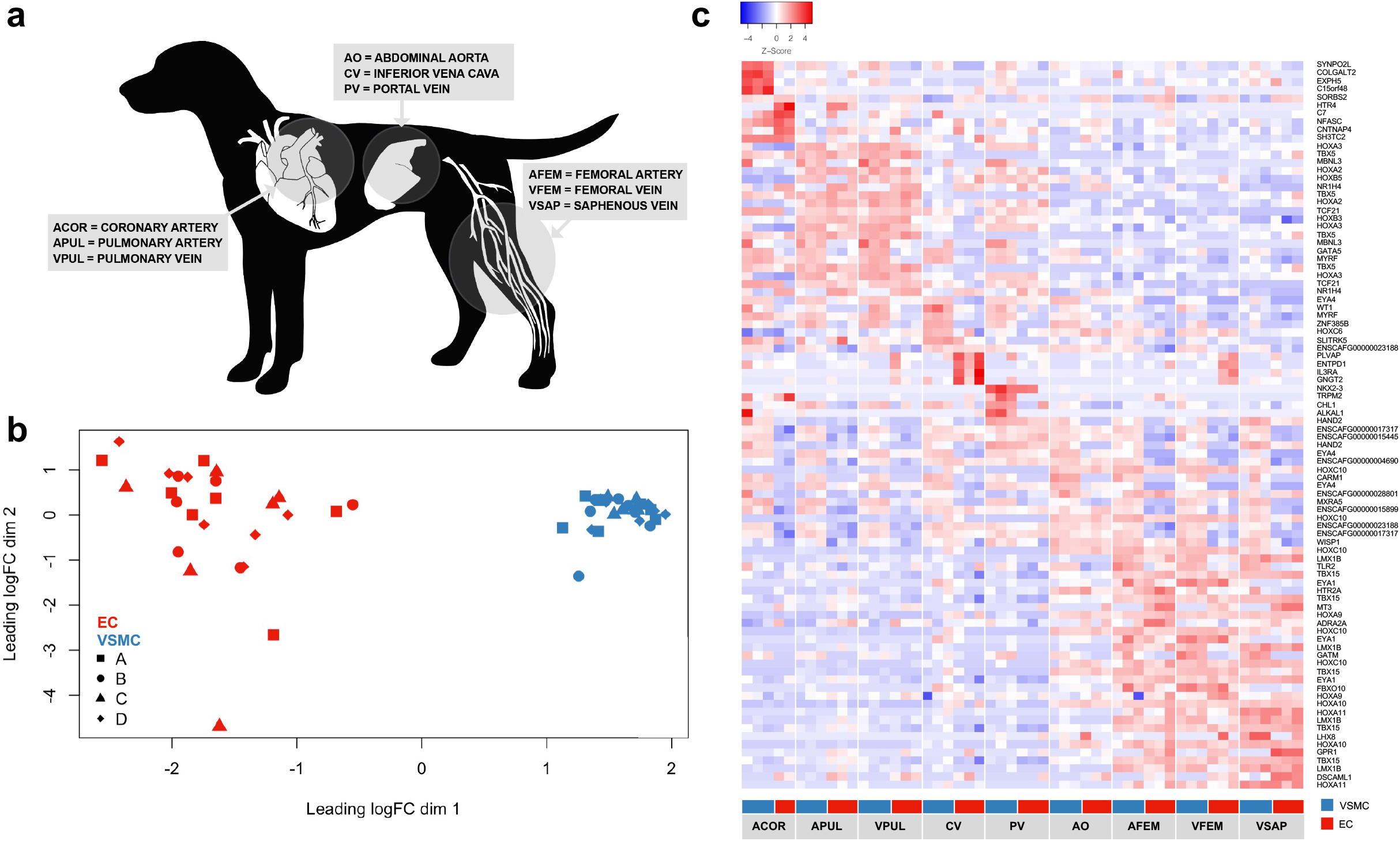
Sample overview. a) Vessel abbreviations and source location of the nine vessels used for endothelial and vascular smooth muscle cell isolation. b) Multidimensional scaling plot of all samples. Colors indicate the cell type and symbols represent the donor (Dog A, B, C or D). VSMCs cluster together on the right side while ECs are more dispersed. There is no apparent donor clustering. c) Heatmap of the top 5 differentially expressed genes per vessel and cell type. Genes are annotated in symbols when available.

Consistent with previous findings, principal component analysis and donor profiles showed distinctive segregation of EC and VSMC cohorts (**Fig. 1b**) which was also confirmed by differential expression analysis as 6,880 genes were found to be significantly up- or down-regulated (*p_adj_* < 0.05) between ECs and VSMCs. The most widely used classification of ECs is based on arterial or venous origin. A differential expression analysis between arterial and venous ECs showed that 258 DEGs are up-regulated in arterial ECs compared to venous ECs and 78 DEGs are up-regulated in venous ECs compared to arterial ECs (**Supplementary Table 1**). No DEGs were identified between arterial and venous VSMCs (**Supplementary Table 2**). Comparing expression profiles based on vessel origin from the nine macro vessels in ECs and VSMCs revealed 3,463 and 2,247 DEGs (*p_adj_* < 0.05), respectively. The entire dataset with the DEGs for each of the EC and VSMC types is provided in **Supplementary Table 3**. The log2 fold change of the DE genes ranged from −7.7 to 6.9 in ECs and from −6.8 to 7.2 in VSMCs, showing substantial variations in gene expression (**Supplementary Table 4**). Groups with the highest number of DEGs are saphenous vein ECs, vena cava ECs, and coronary artery VSMCs with more than 1,000 DEGs each. Less than 20 DEGs were detected in aortic VSMCs, pulmonary artery VSMCs, and femoral artery ECs.

All sample groups show a unique composition of the top DEGs as illustrated by the block-wise pattern in the heatmap build from the top DEGs for each vessel and cell type (**Fig. 1c**). Some overlap in ECs and VSMCs from the same vessel was observed as a total of 194 genes were DEGs in both cell types whereof 14 genes were DEGs in more than one vessel. Highly significant genes (*p_adj_* < 5*10^−7^) expressed in both ECs and VSMCs were *CPQ* and *SH3TC2* for the coronary artery, *HOXC10* for the femoral vein, and *HOXA11*, *HOXA10*, *TBX15*, *LMX1B* for the saphenous vein.

### Functional annotation of differentially expressed genes identifies location-specific vessel processes

Functional annotation showed that several up- or down-regulated DEGs were linked to gene ontology (GO) biological processes or KEGG pathways relevant to the specific location of the vessel (**Supplementary Table 5+6**). Genes regulating the actin cytoskeleton and focal adhesion such as *ITGA7*, *ACTN4*, and *SYNPO2* were expressed at higher levels in arterial ECs. This difference was not observed in VSMCs. More relevant terms were found when the functional annotation analysis was performed using the DEGs per vessel type. Genes involved in heart development and coronary vasculature development including *SMAD6*, *BMP4*, and *NRP1* were found in coronary artery ECs and/or VSMCs. In the up-regulated DEGs of the femoral vein, femoral artery, and saphenous vein ECs and VSMCs, GO terms for biological processes linked to embryonic (limb/skeletal system) morphogenesis were identified. The annotated DEGs in these GO terms contained many homeobox genes as *SHOX2*, *HOXA9*, and *PITX1*.

### Vascular Hox expression in ECs and VSMCs depends on the position of the vessel on the anterior-posterior axis

Next, we hypothesized that homeobox genes represent a group of genes strongly linked to vascular heterogeneity as we observed collective differential expression in proximal vessels. To further investigate this tendency, the quantitative expression of genes containing a homeobox domain was displayed in a heatmap and sorted on sample location (**Fig. 2, Supplementary Fig. 2**). Strong expression of *HOXA1-4* and *HOXB2-6* could be observed for the portal vein, pulmonary artery, and pulmonary vein. HOX genes with high numbers including *HOXA6*, *HOXA7*, *HOXD8*, HOXA9, *HOXA10*, *HOXC10*, *HOXA11*, and *HOXA13* were higher expressed in both cell types for the saphenous vein, femoral artery, femoral vein, and variably enriched for the vena cava and aorta. In the coronary artery *HOXC6*, *HOXC4*, and *HOXA5* were markedly down-regulated compared to the other vessels. In other genes containing a homeobox domain, additional location-specific patterns were observed. In the femoral artery, femoral vein, and saphenous vein *LMX1B*, *IRX2*, *IRX3*, *IRX5*, *ZFHX4*, *EMX2*, *SHOX2*, *TLX2*, *PPRX1* were up-regulated. Cell type patterns were observed for *PRRX2*, *PBX1*, *CUX2*, *PITX2*, *NKX2-5*, *TSHZ3*, *LHX9*, *POU6F1* as these genes were enriched in most VSMCs.

**Figure 2.**
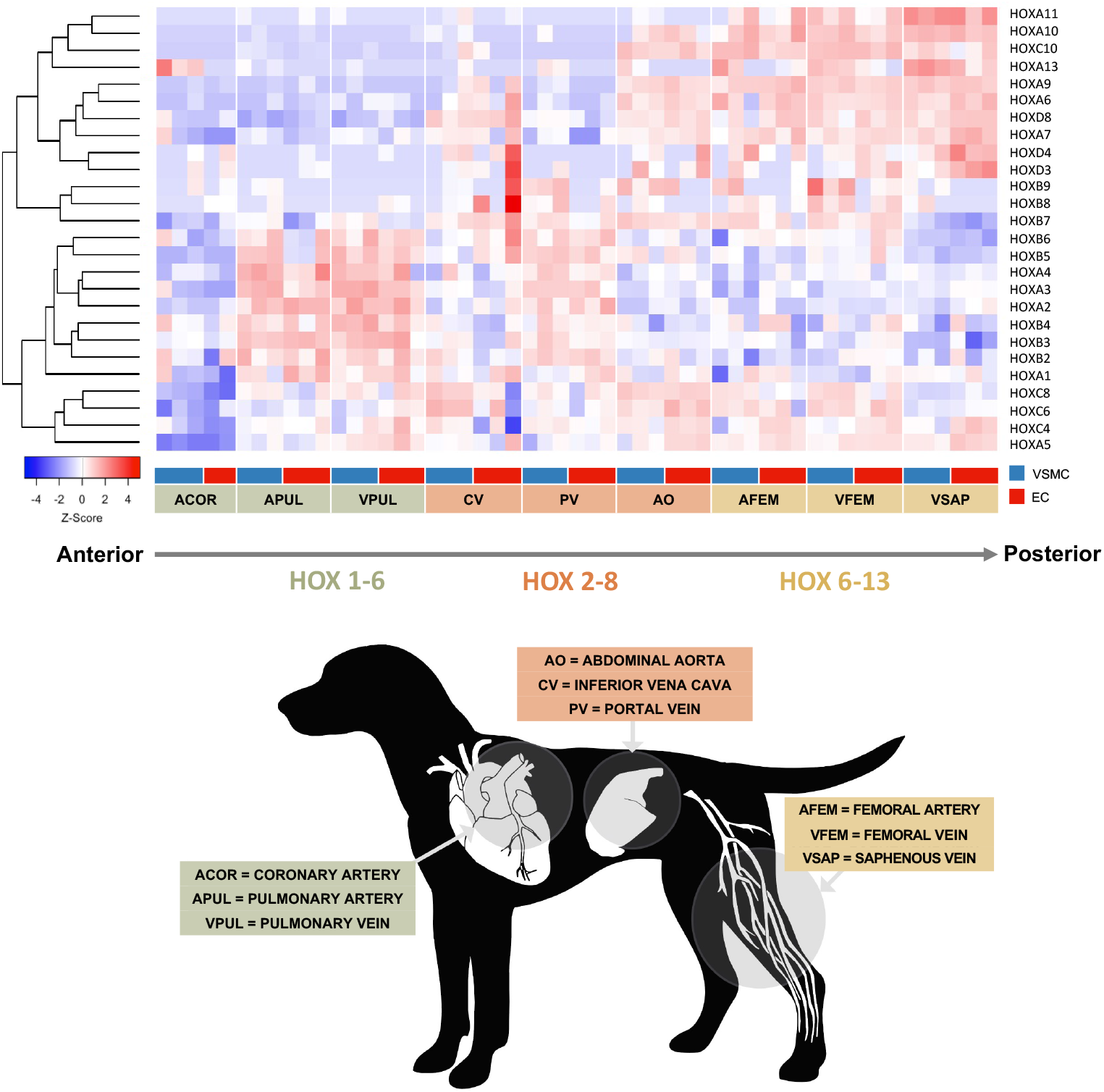
Heatmap of HOX gene expression in endothelial and vascular smooth muscle cells. Vascular Hox expression in ECs and VSMCs depends on the position of the vessel on the anterior-posterior axis, as the expression of Hox genes at the 3’ end (HOX1-6) can be found predominantly in the thoracic vessels and the expression of Hox genes at the 5’ end (HOX6-13) more in the hindlimb vessels. The pattern of the abdominal vessels is less evident.

### Co-expression network analysis identifies several modules related to vascular cell types

The datasets of the EC and VSMC types were used to find groups of co-expressed genes that could be related to the different vessels. The goal of this analysis was to investigate common control mechanisms or activity in related biological processes by studying the correlation between differentially expressed genes with other genes that show a similar pattern across all samples. The weighted co-expression network analysis (WGCNA) is a well-known method that requires a topological overlap matrix to reveal trends of co-expression in the transcriptome. Groups of coexpressed genes are organized in modules and summarized by eigengenes, the typical expression pattern of genes within the module (**Supplementary Table 7+8**). The MDS plot (**Fig. 1b**) showed that cell type is a great source of variation between the expression sets, therefore WGCNA analysis was performed separately for ECs and VSMCs to identify vessel origin as the main driver of heterogeneity. A total of 34 modules for ECs and 31 modules for VSMCs comprising at least 27 genes per module were classified based on hierarchical clustering. Genes not linked to coexpression modules were assigned to M31 (grey module). Subsequently, to ease the comparison of ECs and VSMCs, module names of co-expressed clusters in VSMCs were matched to the most congruent module in ECs (*p* < 0.05). A maximum of one-third of the genes overlapped between the EC and VSMC modules (**Table 1**).

**Table 1.** The number of genes within and overlapping in the modules from the EC and VSMC networks.

|  | EC | VSMC | overlapping genes |
| --- | --- | --- | --- |
| **M3** | 421 | 116 | 36 |
| **M4** | 178 | 137 | 14 |
| **M5** | 380 | 232 | 57 |
| **M6** | 181 | 196 | 14 |
| **M7** | 369 | 357 | 36 |
| **M8** | 408 | 301 | 27 |
| **M9** | 45 | 102 | 3 |
| **M10** | 525 | 520 | 60 |
| **M11** | 90 | 173 | 5 |
| **M12** | 71 | 126 | 6 |
| **M13** | 103 | 133 | 5 |
| **M14** | 112 | 179 | 12 |
| **M15** | 44 | 124 | 19 |
| **M16** | 550 | 1222 | 173 |
| **M17** | 330 | 88 | 5 |
| **M18** | 206 | 212 | 7 |
| **M19** | 1618 | 462 | 217 |
| **M22** | 121 | 371 | 8 |
| **M24** | 145 | 218 | 8 |
| **M26** | 47 | 401 | 4 |
| **M27** | 109 | 127 | 12 |
| **M28** | 409 | 998 | 332 |
| **M29** | 33 | 29 | 14 |
| **M30** | 208 | 266 | 19 |
| **M31** | 8028 | 6822 | 3963 |

### Module construction of co-expressed genes linked to EC subtypes confirms location-specific imprinting

In the EC dataset, 30 of the 34 identified modules were significantly correlated to at least one vessel (**Fig. 3a**). For a selection of modules, the genes with the highest connectivity degree were used to construct hub gene networks (**Fig. 4a**).

**Figure 3.**
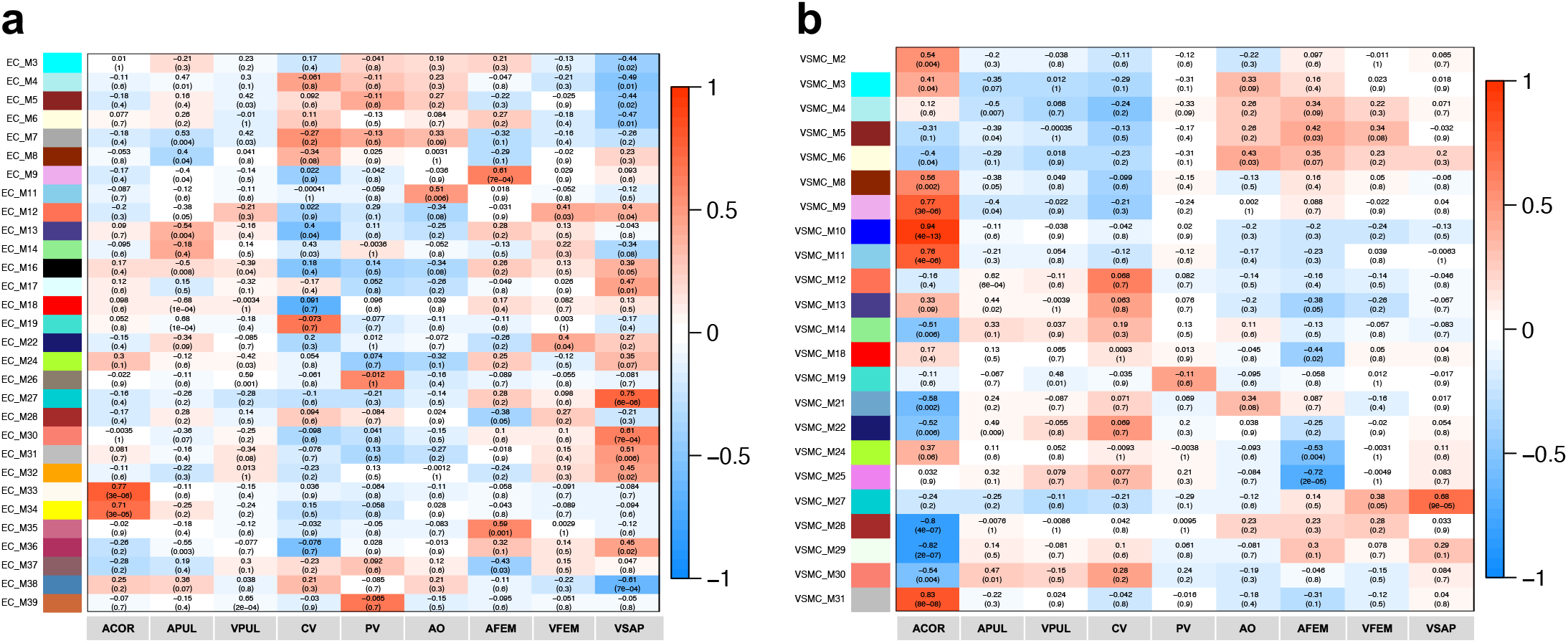
Module-trait relationships of the co-expression analysis in the EC dataset (a) and the VSMC dataset (b). The Pearson correlation coefficient of the module eigengenes (typical expression pattern for genes within the module) and traits (vessel source) was used to calculate the covariance. Each cell of the matrix contains the covariance and the associated p value. A high correlation means that the module eigengene increases with increasing expression in the corresponding sample type.

**Figure 4.**
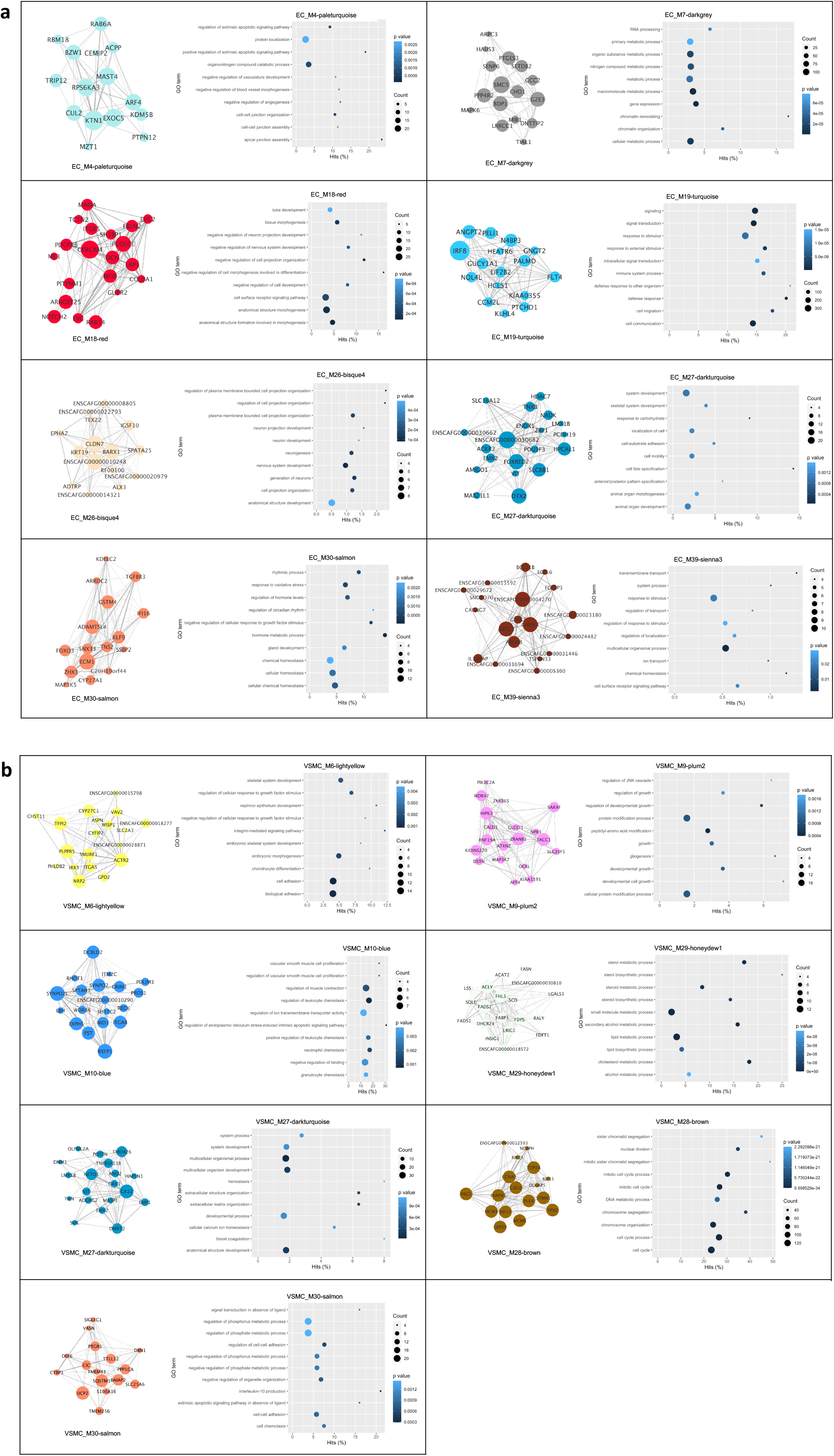
Hub gene networks (left) and Gene Ontology plots (right) in ECs (a) and VSMCs (b). Hub gene networks for a selection of modules that were significantly correlated to one or more vessel types in ECs. For these modules, networks of the top highly connected genes were generated. Node size was scaled to the number of connections with other nodes (degree) and line transparency was dependent on edge weight. The Gene Ontology plots were made of the 10 most significant (non-adjusted p value) biological processes associated with the module. The hits percentage represents the percentage of module genes in the Gene Ontology term. Dot size and color are scaled to the number of module genes in the Gene Ontology term and the p value respectively.

In coronary artery ECs, two very similar modules were significantly upregulated. However, very few genes were annotated due to the relatively small size of these modules. More results were obtained for the vena cava ECs as 9 modules were associated with this sample group. The module with the lowest expression in the vena cava, M18, contained several genes regulating blood vessel development including *ANGPT1*, *EMILIN1*, and *BMPR1A* and genes involved in focal adhesion such as *COL1A2*, *COL6A1-3*, and *PDGFRB*. The largest upregulated module, M19, also contained many genes involved in vascular behavior processes such as endothelial cell migration, angiogenesis, and adhesion. Also, a large number of homeobox genes including *HOXB5-9*, *POU2F1*, *HLX*, and *ZFHX2* were assigned to the upregulated module although most of these homeobox genes were particularly highly expressed in just one of the three samples of the vena cava ECs. Less variance within the sample group with vena cava ECs was seen in the genes assigned to modules M4 and M7. These modules contained, besides genes upregulated in vena cava ECs, also genes associated with aorta and portal vein ECs. These two modules with a similar expression pattern contained the homeobox domain genes *HMBOX1* and *MEOX1* and were linked to basic cellular processes such as cell cycle, differentiation, and response to stress.

In the portal vein ECs, two highly significant homeobox transcription factors *BARX1* and *NKX2-3* were identified as hub genes in module M26 and M39, respectively. Apart from the co-expressed sterol transporters *ABCG5* and *ABCG8* in module M39, only general terms such as metabolism were found for these modules.

In the aorta, pulmonary artery, and pulmonary vein ECs, the accordance between the differential expression analysis and WGCNA was low and no specific ontology terms or pathways were identified.

The best accordance, however, was found in the saphenous vein ECs that shared DEGs with several modules. Module M27 and M30 contained more than 70 percent DEGs for this vessel, and, although not significant, also in the femoral artery and femoral vein ECs a small percentage of DEGs was found. The homeobox transcription factors *HOXA10*, *HOXA11*, *HOXA13*, *HOXC10*, *EMX2*, *LMX1B*, *POU3F3*, *EN1*, *TSHZ1*, and *ZHX3* were present in these modules and many were listed as hub genes (**Fig. 4a**). Genes in this module were annotated to terms like anterior/posterior pattern specification, locomotion, cell adhesion, and cholesterol metabolism.

### Module construction of co-expressed genes linked to VSMCs subtypes confirms location-specific imprinting

In VSMCs, 23 of the 31 modules were significantly correlated with vessel origin (**Fig. 3b**). A large subset of modules was found to be significantly enriched or diminished for the coronary artery VSMCs. From the most positively correlated modules to the coronary artery VSMCs, module M10 contained the highest percentage of DEGs. Several genes were involved in the regulation of the actin cytoskeleton, (vascular smooth) muscle contraction and adhesion such as *SYNPO2*, *NPPB*, *ACTN2*, and *FST* and genes annotated to the cardiovascular system including *TCF21*, *ANGPT1*, and *RECK*. Other positively correlated modules to the coronary artery VSMCs also contained genes annotated to similar processes as represented by hub genes such as *ACTN1*, *ACTA2*, *LEP*, and *HRH2*. The modules downregulated in the coronary artery VSMCs were strongly linked to cell cycle (M28) with hub genes including *MCM3, MCM6, TOP2A*, and *CDC6* and lipid/cholesterol metabolism (M29) with genes like *FADS1*, *ACAT2*, and *FDPS* (**Fig 4b**).

In contrast to the modules in the EC dataset, groups of vessels with a similar enrichment pattern could be distinguished in the VSMCs of the limb and the pulmonary vessels combined with the vena cava. Module M30 is a clear example of a significant positive correlation to the vena cava VSMCs and, although not significant, considerable upregulation in the pulmonary VSMCs. Several genes in this module such as *CX3CL1*, *PDGFRA*, and *HMOX1* were annotated to vascular morphogenesis, cytoskeleton organization, and proliferation. These and other general functional terms were also found in the other modules correlated to vena cava VSMCs.

Analogous to the EC analysis, *BARX1* and *NKX2-3* were among the co-expressed genes enriched for the portal vein VSMCs. In VSMCs however, two additional homeobox domain genes *HHEX* and *LHX6* were also assigned to this module. Genes in this module were significantly linked to the GO terms cell adhesion, defense response, and the regulation of angiogenesis. Some genes including *BARX1*, *RARRES2*, *EDNRB*, and *CTNNB1* were also annotated to the development of the digestive system.

Several modules were collectively enriched for the limb vessels, but module M27 was the only module significantly enriched for the saphenous vein and the femoral vein VSMCs. This module comprising 20 and 9 percent DEGs for saphenous vein and femoral vein VSMCs respectively, contained five genes that were upregulated in all limb VSMCs: *TMEM26*, *LMX1B*, *HOXA10*, and *SLC16A12*. This module contained a total of 7 homeobox genes of which *EMX2* and *LMX1B* were hub genes and *PITX1*, *HOXA10*, *HOXD11*, *EMX2*, *PRRX1* were linked to metabolism. Three other hub genes *CAV1*, *VIT*, and *POSTN* were, together with three other genes in the module, annotated to the extracellular matrix organization.

Module M6, another module with shared DEGs for the limb vessels, aorta, and coronary artery, was significantly correlated to aorta VSMCs and inversely correlated to coronary artery VSMCs. Genes in this module, including the hub gene *ITGA5*, were significantly enriched for focal adhesion the regulation of actin cytoskeleton. Besides the homeobox genes *HOXA9*, *HOXC10*, and *SHOX2* that were annotated to skeletal system development, three other homeobox genes namely *IRX3*, *IRX5*, and *ZEB1* were included in this module.

## 3 Discussion

In our prior work we documented the presence of diversity in EC and VSMC subtypes.^6, 15^ In this study, we adopted a simple and rapid method for isolating and characterizing ECs and VSMCs from the same vessel to identify the persistent vessel and cell type-specific expression patterns that define the basis of vascular heterogeneity in cultured canine ECs and VSMCs upon removal from the natural environment.

Among the differentially expressed genes, a noticeable number of genes with a role in embryonic development were identified. These genes may be the basis of positional memory because once cell fate is determined, expression of some of these genes was shown to be maintained in adult tissues to regulate regeneration in response to injury.^16^ Key players in the positional memory of many cell types including ECs and VSMCs are the Hox genes, a subset of the homeobox domain genes.^17, 18^ Hox genes, the most well-characterized group of homeobox genes, are known for their collinearity (correspondence between the sequence of Hox genes along the chromosome and their expression along the anterior-posterior axis) during embryonic development in all bilateral animals.^19^ In line with this, our analyses suggested that vascular Hox expression in ECs and VSMCs depends on the position of the vessel on the anterior-posterior axis, as the expression of Hox genes at the 3’ end (*HOX1-6*) was found predominantly in the thoracic vessels and the expression of Hox genes at the 5’ end (*HOX6-13*) more in the hindlimb vessels.^19^ Vascular spatial collinearity has been previously described in several endothelial and supporting vascular cell types in different species.^17, 18, 20, 21^ Our results thus lend further credence to the earlier findings that one may expect to find clustering of Hox genes analogous to the position of the vessel on the body axis. The function of Hox genes in the adult vasculature has not been entirely elucidated although some studies suggest a role for Hox in angiogenesis and vascular remodeling.^22–24^

We also identified expression of other embryonal transcription factors and homeobox domain genes that do not belong to the subgroup of Hox genes. For example, our data showed that *LMX1B* (associated with hindlimb ECs and VSMCs) was co-expressed with *EMX2*, a known target of *LMX1B* during limb development.^25^ *TBX15*, another gene differentially enriched for all limb samples, regulates the formation of the skeleton elements of the limb, vertebral column, and head.^26^ Their functional impact on vascular homeostasis is yet to be discovered but as several genes in the co-expression module were associated with extracellular matrix formation in VSMCs and cholesterol metabolism in ECs, these genes may play a role in these processes as well.

Coronary artery VSMCs displayed a highly divergent expression pattern compared to the other VSMC types. The proepicardial origin of these VSMCs may, at least in part, account for this difference. In line with this, the co-expressed genes *CRIM1* and *TBX18*, known to be highly expressed in the proepicardial organ, were found to have sustained transcription in coronary artery VSMCs in our study.^27–29^ Since a disturbed lipid metabolism is a key element of atherosclerosis, the discovery of a group containing genes involved in lipid metabolism that was strongly negatively associated with coronary artery VSMCs, attracted our attention.^30^ However, lipid metabolism is also closely related to the differentiation state.^31^ Other indications for an altered differentiation state in coronary VSMCs are provided by the significant overexpression of contractile markers such as *ACTA2*, *ACNT1*, and *MYL9* and the strong down-regulation of co-expressed genes involved in cell cycle regulation suggesting a lower proliferation rate or even cellular senescence in these samples.^4, 32, 33^

For the pulmonary artery and vein several highly differential expressed genes were identified. For example, *TBX5*, a gene is crucial in cardiac morphogenesis and lung development, was enriched for the pulmonary vessels and the coronary artery.^34, 35^ *TCF21*, enriched for the portal vein, pulmonary artery, and pulmonary vein in ECs and the coronary artery in VSMCs, has also been found to be expressed in tissue contributing to the developing lung, heart, and gut.^36^

In the positional memory of portal vein samples, we identified several genes involved in gastrointestinal development. The DEGs for portal vein ECs and VSMCs *HAND2, BARX1*, and its intermediate transcription factor *ISL1* have been linked to gut rotation and gastrointestinal functioning.^37–39^

Vena cava VSMCs share a similar co-expression pattern with pulmonary VSMCs while vena cava ECs are more similar to the aorta and the portal vein. The comparable co-expression patterns may be explained by the mesothelial origin of both vena cava and pulmonary VSMCs. Lineage tracing of *WT1* (the number one DEG in our vena cava VSMCs) in mesothelial cells showed that the majority of the VSMCs in the gut and approximately one-third of the VSMCs in the lung vasculature had a mesothelial origin.^40, 41^

Among the vena cava EC samples, we detected one outlier that expressed several typical EC markers such as *VWF* and *CD34* at very high levels. Since the other two vena cava samples expressed these genes at levels comparable to the entire set of ECs, these results are probably not representative of the positional memory of vena cava ECs but rather a result of fewer divisions due to a higher yield after isolation. In line with observations by others, these endothelial markers most likely faded out in the other EC samples during culture.^7^

To conclude, this study shows that adult vascular cells in culture express a remarkably high amount of transcription factors crucial to organ development in the embryo. The persistent expression of these genes in culture indicates that these genes are not regulated by flow or surrounding cell types but are rather fixed in the molecular memory. The identification of genes that are part of the positional memory of vascular cell types provides us with better tools for tissue transplantation, genetic (re)programming, and regeneration. Matching this basic expression profile comprising Hox genes and other embryonic transcription factors to the site of transplanted tissue, for example, may increase the success rate of tissue implementation. This study shows that we are still at the beginning of understanding angiodiversity. Expanding the data collection may reveal additional genes that are important to the key position of the cardiovascular system in homeostasis and disease.

## 4 Methods

### Ethics statement

For this study, blood vessels from healthy male dogs (*n*= 4; age 12–14 months, with an average of 26 kg) were collected. The vessels were obtained as surplus material from fresh canine cadavers used in unrelated research on pacemakers which was performed according to the Dutch Experiments on Animals Act and conform to the EU standards.^42^ Acquiring this tissue did not influence the method and moment of euthanasia.

### Whole transcriptome sequencing of vascular cells

ECs and VSMCs were isolated from the aorta, coronary artery, caudal vena cava, vena porta, femoral artery, femoral vein, saphenous vein, pulmonary vein, and pulmonary artery (Fig. 1A) according to a previously published protocol.^6^ The isolation of these two different cell types was confirmed on gene and protein expression, and morphological and functional levels.^15^ In short, a vessel of approximately 5 cm in length was inverted inside out to bring the endothelial cell layer to the outside of the vessel. Both ends of the vessel were closed by placing purse string sutures to prevent exposure of the non-endothelial vascular tissue to the digestion medium consisting of collagenase II (0.15 U/mL, Life Technologies) and dispase (0.15 U/mL, Life Technologies) in Dulbecco’s Modified Eagle’s Medium (DMEM) GlutaMAX (Life Technologies). After each vessel was digested for one hour, the vessel was removed and the cell suspension was centrifuged at 1500 rpm for 5 minutes. The cell pellet consisting of ECs was suspended in Canine Endothelial Growth Medium (CECGM, Cell Applications, San Diego, CA, United States) and added to gelatin (Sigma–Aldrich, Saint Louis, MO, United States) coated 6-wells plates. To isolate the VSMCs, the remaining tissue was chopped finely and digested with collagenase type II (0.09 U/mL) in DMEM GlutaMAX. After 4 hours, a single cell suspension was obtained by filtration through a 70 μM strainer. The cell suspension was centrifuged at 1500 rpm for 5 minutes and resolved in a culture medium consisting of DMEM glutamax, 10% Fetal Calf Serum (FCS, Life Technologies), and 100 μg/mL Primocin (Invivogen, Toulouse, France) before adding the cells to the culture plates.

All cells were maintained in a humidified incubator with 5% CO^2^ at 37 °C and passaged when 70-80% confluence was reached.

Per vessel and cell type, three samples (*n*=3) were selected. RNA was extracted from a cell pellet with approximately one million ECs or VSMCs at the third passage using an RNeasy Mini Kit (Qiagen, Venlo, Netherlands). The remaining genomic DNA traces were removed with an on-column DNase digestion (Qiagen, Venlo, Netherlands).

Poly(A) Beads (NEXTflex) were used to isolate polyadenylated mRNA, from which sequencing libraries were made using the Rapid Directional RNA-seq kit (NEXTflex). Libraries were sequenced using the Nextseq500 platform (Illumina), producing single-end reads of 75bp. Reads were aligned to the canine reference genome CanFam3.1 using STAR version 2.4.2a (https://github.com/UMCUGenetics/RNASeq).

The raw and analyzed files have been uploaded to Gene Expression Omnibus under accession number GSE171437.

### Differential expression analysis

The RNA-sequencing data was merged with the publically available dataset GSE118029^15^ and analyzed by the workflow outlined by Law et al.^43^ Briefly, the dataset, consisting of 56 samples, was imported, organized, filtered, and normalized for composition bias using trimmed mean of M-values (TMM) in the edgeR package.^44^ Data quality was assessed by including three technical replicates which were excluded from further analysis after testing for reproducibility (Spearman R^2^□>□0.9). In our analyses, counts per million (CPM) and log2-counts per million (log-CPM) transformations were applied to account for library size differences. Genes were filtered on worthwhile counts by using the filterByExpr function that filters genes based on the median library size and the number of samples per group. The limma package was used to apply the voom method which estimates the mean-variance relationship of log-counts, generates a precision weight for each observation and enters these into the limma empirical Bayes moderation pipeline to assess differential gene expression.^45, 46^ Differential expression was calculated for contrasts across all sample groups within the same cell type. The model matrix was processed by *contrast.fit* before being passed to *eBayes* in the Limma package which resulted in an adjusted p value and a logarithm-transformed fold change (logFC) for each contrast. A multidimensional scaling plot was generated from 11,992 genes.

### Functional enrichment

Gene annotations were obtained from the Ensembl BioMart database.^47–49^ Gene set enrichment analysis (GSEA) was performed using the GOseq Bioconductor package and the enrichKEGG function of the clusterProfiler package.^50, 51^

### Co-expression network construction

Preprocessing of the datasets was performed by the variance stabilizing transformation as implemented by the package DESeq2.^52^ Co-expression was calculated using the Weighted Gene Co-expression Analysis (WGCNA) R package.^53, 54^ Network construction with the WGCNA algorithm entails the calculation of an adjacency matrix by raising the connectivity between genes to a soft thresholding power. The optimal soft thresholding power was chosen using the scale-free topology criterion (**Supplementary Fig. 1**). Subsequently, a signed network resulting in the separation of inversely correlated genes to different modules was constructed. Using a topological overlap matrix, network interconnectedness required for hierarchical clustering was calculated. For the detection of modules in the cluster tree using the *cutreeHybrid* function, the minimum module size was set on 27, and the detection sensitivity on 1.^55^ Modules with similar eigengenes were merged, as measured by a correlation of eigengenes above 0.8.

### Functional annotation

The correlations of the module eigengenes with vessel origin were calculated and displayed in a module-trait heatmap. Modules without significant correlation with any vessel were excluded from this plot. Functional annotation was performed through the utilization of the GOseq function in the Bioconductor package and the enrichKEGG function of the clusterProfiler package on genes assigned to each module.^50, 51^ For all functional terms with more than 3 genes per terms, the adjusted p value was calculated using the Benjamini and Hochberg false discovery rate.^56^ Besides significance, relevant gene ontology terms and KEGG pathways were selected based on the number of annotated genes and their role in vascular functioning.

### Hub gene selection and visualization

Hub genes are the most influential genes within a module and therefore possibly exert strong biological importance. In this study, intramodular hub genes are defined as genes with both a high correlation to the module eigengene and a high number of connections with other genes in the module. Firstly, genes with a membership value above 0.7 were collected as potential hub genes. Secondly, using Cytoscape (Version 3.7.2) with its plugin NetworkAnalyzer, the number of edges between these genes was extracted from the topological overlap matrix.^57–59^ To obtain consistent results from all modules, the average adjacency value of the selected genes was used as a cut-off threshold for the edges. Finally, intramodular hub genes were ranked on the number of connections as visualized by their node size in the Cytoscape networks.

## 5 Acknowledgments

This research was partially funded by the K.F. Hein Fund and the ERA-CVD SCALE project (2019T109). The authors thank the Utrecht Sequencing Facility (USF) for their support in the whole-transcriptome sequencing, partially subsidized.

## 6 Contributions

F.G.S. conceived the idea, H.S.K., B.S. and L.A.O. helped to design the study. H.S.K., L.A.O and T.C.F.S. collected the tissues. L.A.O and T.C.F.S. cultured the cells and isolated the RNA. M.M. and N.A.M.D. sequenced the samples. T.C.F.S. analyzed the data with the help of M.H. and F.G.S.. T.C.F.S. wrote the manuscript with critical inputs from M.H., M.M., L.A.O., B.S. and F.G.S..

## 7 Competing interests

The authors declare no competing interests.

## Supplementary tables are available upon request

Supplementary Table 1. Differential expression analysis of arterial versus venous endothelial cells (ECs). Positive log fold changes (logFC) indicate higher expression in arterial ECs while the negative values represent a higher expression in venous ECs.

Supplementary Table 2. Differential expression analysis of arterial versus venous vascular smooth muscle cells (VSMCs). Positive log fold changes (logFC) indicate higher expression in arterial VSMCs while the negative values represent a higher expression in venous VSMCs.

Supplementary Table 3. Matrix of significantly differential expressed genes (*p_adj_* < 0.05) in all sample groups. Upregulated gene = 1, down-regulated gene = −1.

Supplementary Table 4. Top 5 differentially expressed genes per vessel and cell type. Genes are annotated in symbols when available. For each gene, log2-fold-change (LogFC), average log2-expression (AveExpr), moderated t-statistic (t), raw p value (*p*), adjusted p value (*p_adj_*), and log-odds that the gene is differentially expressed (B) were calculated.

Supplementary Table 5. Gene ontology terms for contrasts of all sample groups.

Supplementary Table 6. KEGG pathways for contrasts of all sample groups.

Supplementary Table 7. WGCNA analysis endothelial cells

Supplementary Table 8. WGCNA analysis vascular smooth muscle cells

---

**Supplementary Figure 1.**
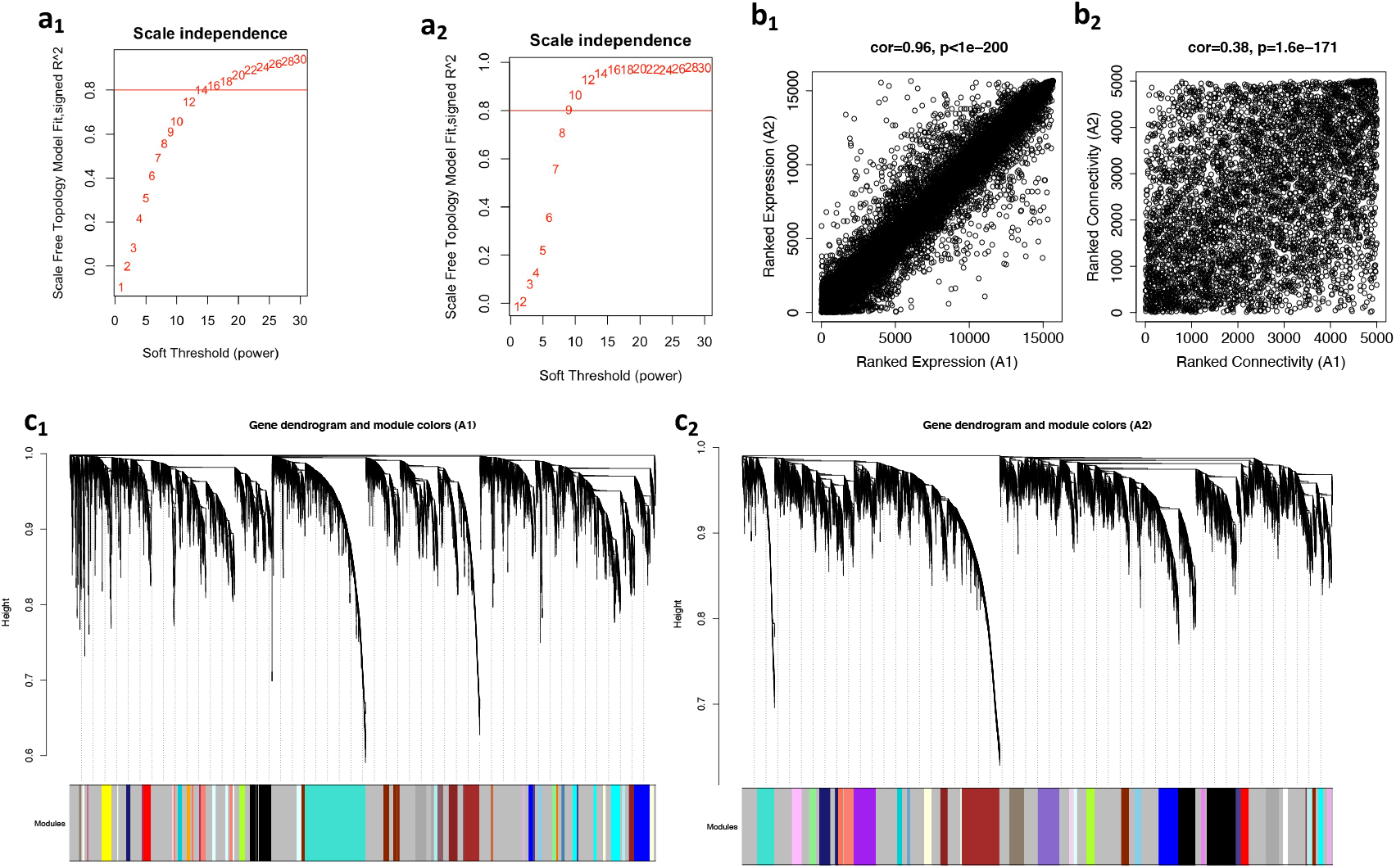
Statistics used for the weighted gene co-expression analysis. a_1&2_) A scale free topology fit index was made to determine the optimal value for the soft threshold for the EC dataset (a_1_) and the VSMC dataset (a_2_). Numbers in the plots illustrate the soft thresholding powers. The approximate scale-free topology can be achieved for ECs at the soft-thresholding power of 14 and for VSMCs of 9. b_1&2_) The correlations between the average gene expression (b_1_) and overall connectivity (b_2_) between the EC (x-axis) and VSMC (y-axis) datasets used to assess the comparability. c_1&2_) A gene dendogram based on the topological overlap dissimilarity index showing the modules identified by WGCNA in the EC (c_1_) and VSMC (c_2_) dataset.

**Supplementary Figure 2.**
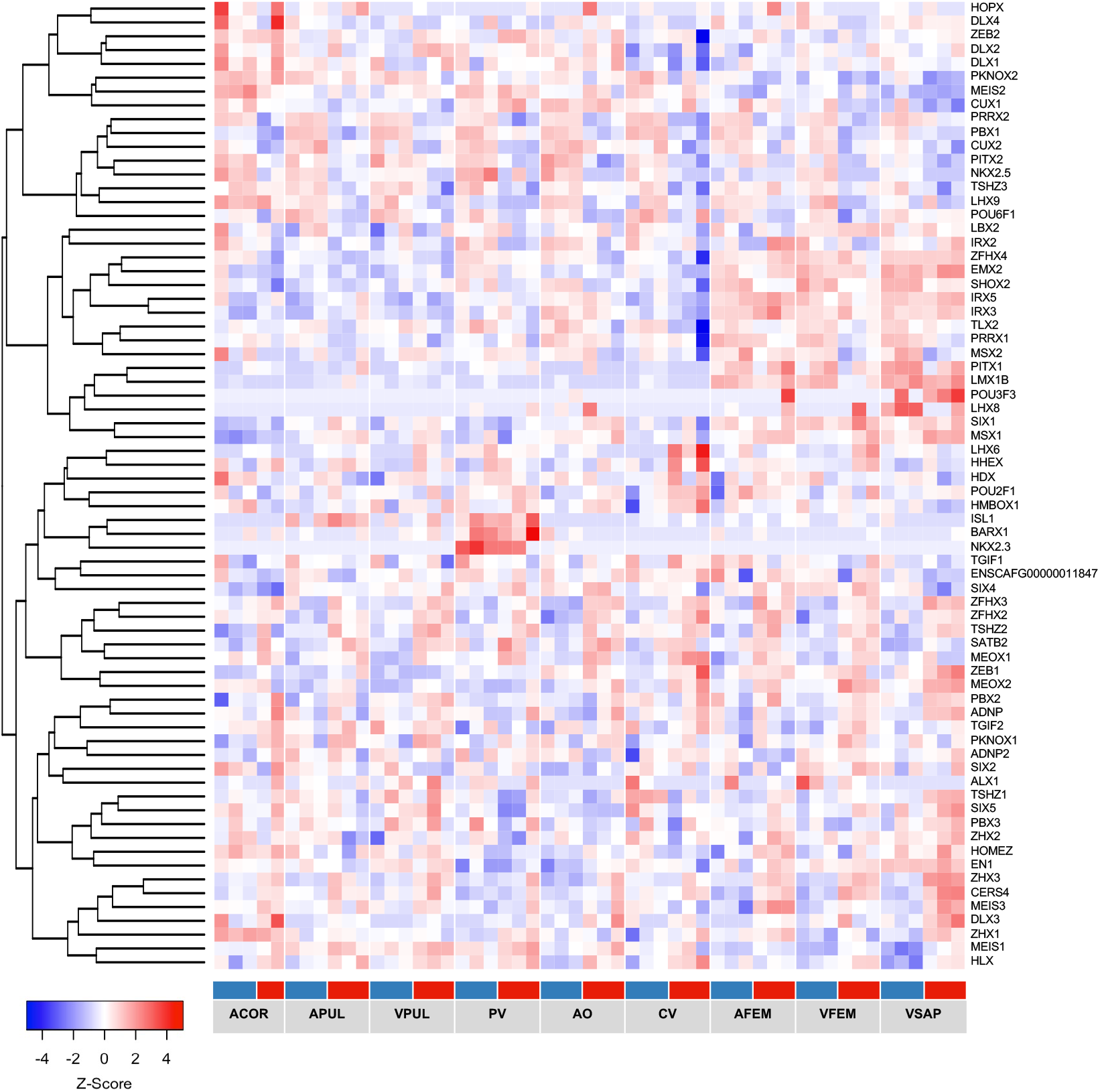
Heatmap of genes with a homeobox domain (SMART SM00389, other than HOX) with expression in ECs and VSMCs.

## Notes

### Competing Interest Statement

The authors have declared no competing interest.

